# The genetic landscape of animal behavior

**DOI:** 10.1101/139246

**Authors:** Ryan A York

**Affiliations:** Department of Biology, Stanford University, Stanford CA, 94305, USA

## Abstract

Although most animal behaviors are associated with some form of heritable genetic variation we do not yet understand how genes sculpt behavior across evolution, either directly or indirectly. To address this, I here compile a dataset comprised of over 1,000 genomic loci representing a spectrum of behavioral variation across animal taxa. Comparative analyses reveal that courtship and feeding behaviors are associated with genomic regions of significantly greater effect than other traits, on average three fold greater than other behaviors. Investigations of whole-genome sequencing and phenotypic data for 87 behavioral traits from the Drosophila Genetics Reference Panel indicate that courtship and feeding behaviors have significantly greater genetic contributions and that, in general, behavioral traits overlap little in individual base pairs but increasingly interact at the levels of genes and traits. These results provide evidence that different types of behavior are associated with variable genetic bases and suggest that, across animal evolution, the genetic landscape of behavior is more rugged, yet predictable, than previously thought.

## Introduction

Nearly all behaviors are associated with some form of heritable genetic variation (Kendler and Greenspan 2006). This interplay between genetic and other forces that shape behavior is complex and disentangling it occupies an array of research endeavors, spanning disciplines from evolutionary biology to psychiatry. Accordingly, recent years have seen reasonable progress toward understanding the genetic architecture of certain behavioral traits using model systems (Reaume and Sokolowski 2011). The general conclusion from this research in mice, flies, worms, and humans is that the genetic architectures of behaviors generally fit an exponential distribution, with a small number of loci of moderate to large effect and a larger number of loci with small effects (Robertson 1967; Flint and Mackay 2009). However, owing to limits in data and methods, the extent to which genetic architectures vary across a full spectrum of behaviors and animal taxa has remained largely unexplored.

Behaviors can exhibit considerable variation in genetic influence. Comparative analyses reveal that behaviors vary substantially in heritability estimates, most often ranging between 10% and 50% (Kendler and Greenspan 2006; Mousseau and Roff 1987; Meffert et al. 2002). Analyses of individual behaviors reveal even greater diversity. For example, a single retro-element is responsible variation in a courtship song between *Drosophila* species (Ding et al. 2016) while other traits, such as deer mouse burrowing, have modular genetic architectures comprised of multiple interacting loci (Weber et al. 2013). Furthermore, the structure and effect of genetic architectures may vary with behavioral traits, as suggested by the preponderance of large effect loci found for insect courtship traits across multiple species (Arbuthnott 2009). Despite these observations the extent to which behavioral traits may systematically vary across species and behaviors remains unknown. Understanding this could provide insights into how behaviors respond to evolutionary processes, the prospects for finding general principles in the genetic evolution of behavior, and even potentially why there has been such variable success in the mapping of human neuropsychiatric traits.

Here, using reports associating behavioral variation with the genes for specific traits across diverse species, I assemble a comparative behavior genetics resource composed of 1,007 significant genomic loci from 114 QTL studies conducted in 30 species across 5 taxonomic classes. These data exploit advances in sequencing and genetic marker design that have accelerated reports using quantitative trait locus (QTL) mapping to identify genomic regions that are associated with behavioral variation (Lander and Botstein 1989; Flint and Mackay 2009). With the compiled dataset I compare the genetic architecture of behavioral types across animal taxa. I then corroborate these observations and assay genetic processes involved in the early stages of behavioral differentiation in a natural population using whole genome data from the Drosophila Genetic Resource Panel (DGRP). These analyses provide insight into the genetic architecture of behavior across animals and the interplay between specific behavioral traits and their genetic influence through evolutionary history.

## Results and Discussion

I performed a comprehensive analysis of results aggregated from 114 QTL studies conducted in 30 species across 5 taxonomic classes to assemble a comparative behavior genetics resource composed of 1,007 significant genomic loci (Database S1). The species examined represent over 500 million years of evolutionary divergence and over a broad spectrum of phylogenetic data (Fig 1a). For each locus I annotated the trait measured and its associated effect size (percent phenotypic variation explained), the reported measure of significance (e.g., LOD score), genomic locus, and study sample size. I focused the analyses on the reported effect sizes to allow comparison of the genomic architecture of traits across studies similar to previous meta-analyses of behavioral QTL in mice and flies (Flint 2003; Flint and Mackay 2009).

**Fig. 1.**
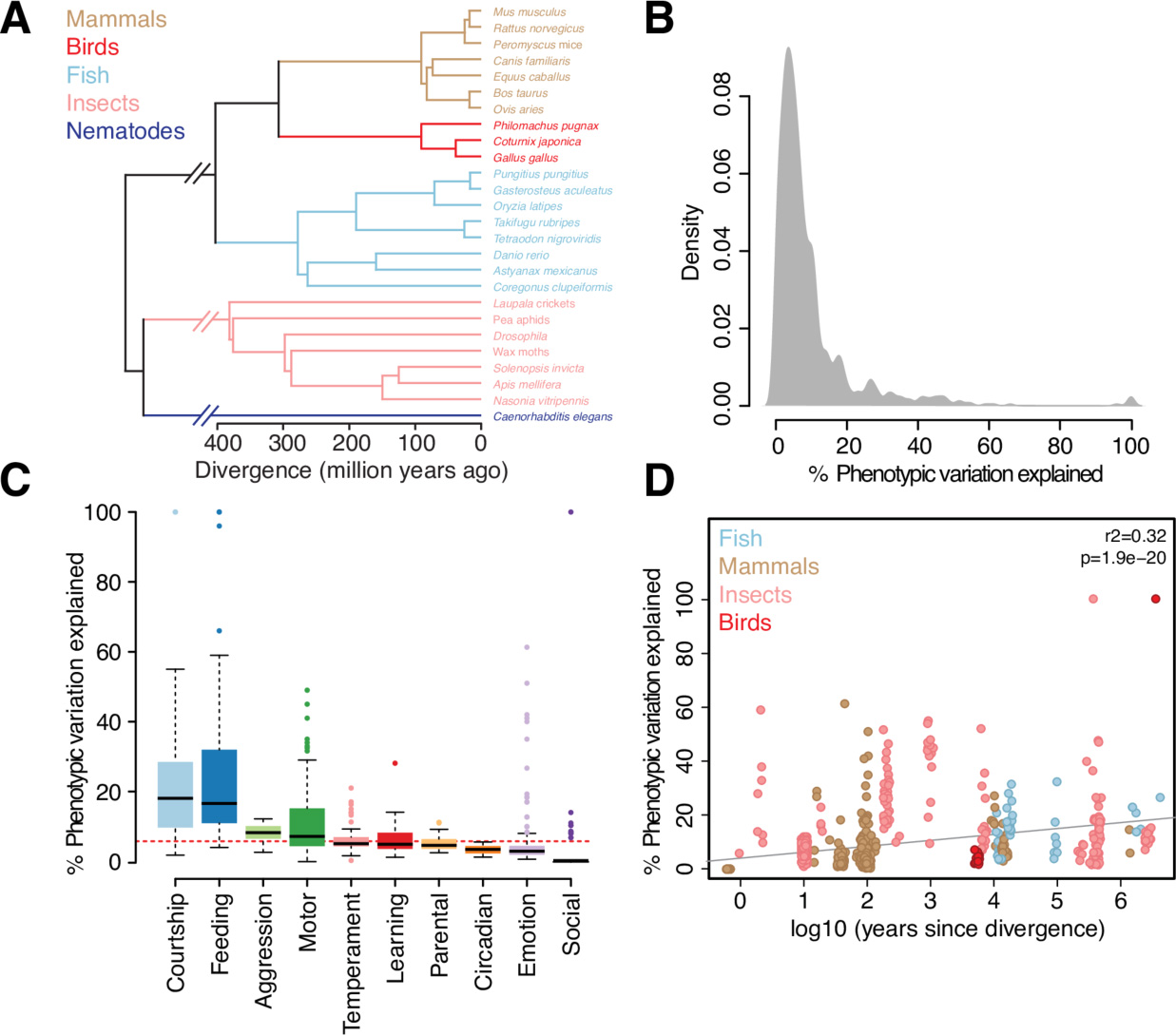
The genomic landscape of animal behavior. **(A)** Phylogeny of all species studied in which genomic loci were collected for the meta-analysis. **(B)** Density plot of the distribution of effect sizes for all behavioral traits studied. **(C)** Boxplot of effect sizes (% variation explained) by behavioral category. **(D)** Scatterplot of the relationship between evolutionary divergence (represented by the log10 of years since divergence) and effect size.

I found that the distribution of effect sizes in the dataset is similar to that found in these previous studies (Fig 1b). In the majority of loci (89.51%) the effect sizes are less than 20% with a mean effect size of 9.54%, suggesting that the genetic bases of most behaviors assayed are complex and composed of many loci of moderate effect.

Though these results support a model of many loci with small effects for behavior overall, I then asked whether genetic architecture might vary across *types* of behavior. I identified ten behavioral categories for which traits had been measured in at least two species (See supplementary methods). My null hypothesis was that individual categories would likely reflect the overall distribution seen across the dataset, consistent with previous observations that QTL have relatively similar effect sizes across mouse and fly phenotypes (Flint 2003; Flint and Mackay 2009). Surprisingly, I found instead that behaviors differed significantly in their effect sizes. Specifically, loci associated with courtship (n=124) explained significantly more phenotypic variance than all other behaviors combined (Kruskal-Wallis p = 6.7 × 10^−29^) and had a mean effect size three times larger than found in all other categories (Fig. 1c). Loci associated with feeding behaviors (n=11) also explained significantly more phenotypic variance than all other behaviors combined (p = 6.8×10^−13^) while emotion and social behaviors explained significantly less (p = 8.6 × 10^−33^; p = 2.5 × 10^−21^, respectively). These data suggest that, across species, courtship and feeding behaviors possess genetic architectures different from those of other traits.

To assess whether these observations arose from differences in the behavioral traits, as byproducts or experimental artifacts I controlled for factors that might have contributed bias. I first considered the effect of *intraspecific* (within species) compared to *inter*specific (between species) crosses used for the QTL mapping, a known source of influence in QTL studies (Broman 2001). I indeed found that experiments employing interspecific crosses identified loci of significantly higher effect (p = 4.5 × 10^−5^). To control for this quantitatively, I estimated phylogenetic divergence and generation times between the crosses used in each of the 115 studies (Supplementary methods). There was a positive correlation between evolutionary divergence and effect size (r^2^ = 0.32, p = 1.9 × 10^−20^; Fig. 1d; Supplementary methods). I also considered sample size, a well-known source of bias for which, as might be expected, there was a negative correlation with effect size (r^2^ = −0.37, p < 0.0001).

To test the effect of key variables, evolutionary divergence individually, sample size individually, and both combined, I used three linear models (Supplementary methods). Strikingly, the overall structure of the effect size distribution remained largely unaffected after analysis of the residuals from all three models (Fig S1-3; Supplementary methods). In addition, courtship and feeding behaviors had significantly larger effect sizes even after accounting for these potential sources of bias (p = 1.4 × 10^−14^ and p = 5.7 × 10^−7^, respectively; Fig 2a).

**Fig. 2.**
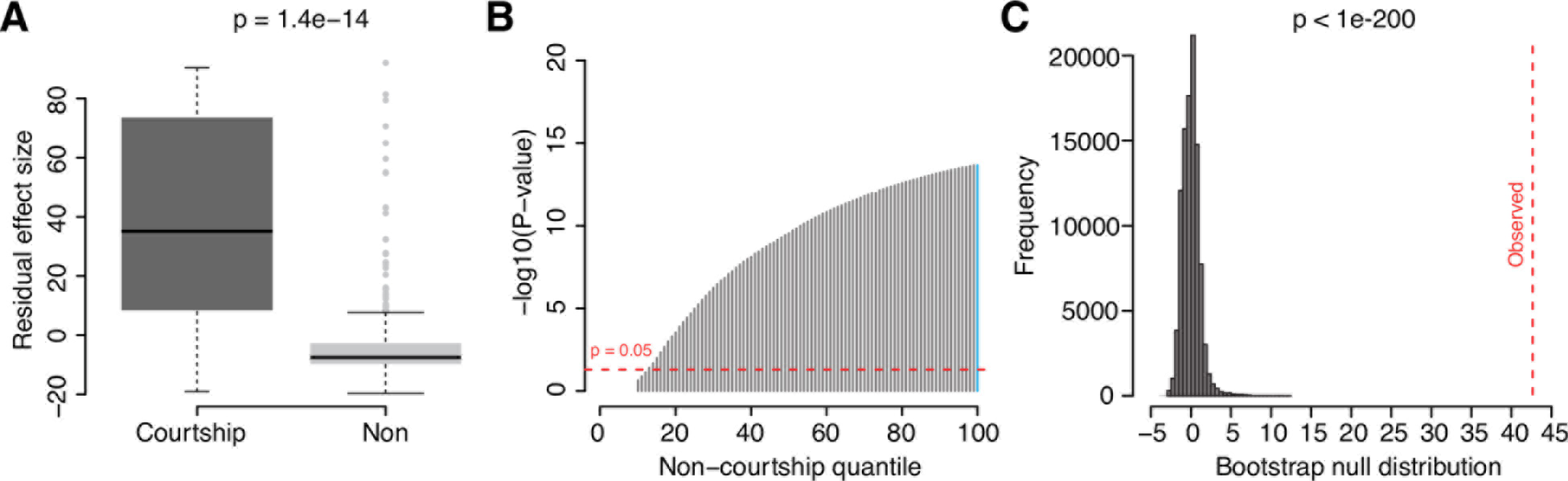
Assaying the genetic architecture of courtship. **(A)** Boxplot of the comparison between the residual effect sizes of courtship and non-courtship behaviors that resulted from a linear model controlling for sample size and evolutionary divergence. **(B)** Quantile-based –log10(p-values) comparing the residual effect sizes of courtship and non-courtship behaviors at each quantile cutoff. The blue line corresponds to the comparison in Fig 2a. **(C)** The null distribution resulting from bootstrapping all non-courtship residual effect sizes for 10,000 permutations and the observed median residual effect size for courtship (dashed red line).

After eliminating sources of potential biases inherent to individual datasets, I next considered the possibility that the detection of courtship and feeding behaviors as outliers was a trivial outcome of our own classification method for grouping single behaviors into ten categories. Minimally assuming that the categorizations of courtship and feeding traits were correct, it is possible that the binning of traits into the other eight categories may have masked a real signal from some biologically relevant categorization.

To test this possibility, I compared the distribution of effect sizes for the courtship and feeding categories to the distribution for *all* other behaviors combined (Supplementary methods). I found that courtship behaviors explained significantly more variation (p <0.05) than 89% of non-courtship behaviors while feeding behaviors explained more variation than 46% of non-feeding behaviors (Fig 2b; Fig S4b). I complemented this test with a bootstrap analysis that created a null distribution from 10,000 permutations of the non-courtship/feeding trait effect sizes. The observed mean adjusted effect size for both courtship and feeding fell significantly outside the bootstrap null distribution created for each comparison (p < 5 × 10^−200^)(Fig 2c; Fig S4c). These findings reject the notion that there may be another categorization of non-courtship and feeding behaviors missed by our schema that explains substantially more variation of effect.

My results suggest that courtship behaviors, and to a lesser extent feeding, may respond to evolutionary pressures differently than other behavioral traits. Consistent with this notion, previous analyses of the QTL behavior literature in insects found that a majority of courtship traits are associated with few loci of particularly strong effect that play a potential role in rapid speciation through prezygotic isolation (Arbuthnott 2009). In addition, theoretical work has suggested that traits controlling local adaptation during speciation, such as courtship and feeding, evolve more rapidly if they are associated with a smaller number of loci (Gavrilets et al. 2007). Given the importance of behavior’s role in the early stages of speciation it may be possible that for the organisms and traits analyzed here, courtship and feeding traits with simpler genetic components of large effect were selected for during the evolution of these lineages. These observations led me to hypothesize that, in a naturally interbreeding population, courtship and feeding behaviors may be associated with more heritable genetic architectures of greater effect when compared to other behavioral traits.

To test this idea, I used the Drosophila Genetic Reference Panel (DGRP). The DGRP is comprised of over 200 inbred, fully sequenced *Drosophila melanogaster* lines isolated from a farmer’s market in Raleigh, North Carolina (Mackay et al. 2012). Phenotypic measures for a wide number of behavioral traits are available for the DGRP lines in addition to full genome sequence and variant information, making this resource unique in enabling us to ask larger scale questions about variation and evolution in behavior. I collected phenotypic measures for 87 behavioral traits spanning 8 categories, produced in 9 separate GWA studies (Jordan et al. 2012; Weber et al. 2012; Swarup et al. 2013; Arya et al. 2015; Gaertner et al. 2015; Garlapow et al. 2015; Morozova et al. 2015; Shorter et al. 2015).

I first used genome-wide complex trait analysis (GCTA) to survey the extent to which the 87 behavioral traits varied in genomic heritability attributable to all autosomal SNPs (Yang et al. 2011). After running GCTA 20 behavioral traits passed a p-value threshold of 0.05, indicating that autosomal SNPs could explain more trait variation than by chance in these cases (Fig. 3a; Supplementary methods). The majority of these traits were enriched for involvement in courtship and feeding: 30% (6/20) were associated with courtship and 50% (10/20) were either involved in olfactory behavior or feeding. Notably, for a number of these traits the vast majority of phenotypic variation could be explained by genome-wide SNPs, including preference for the food odorant ethyl acetate (99.99 +/−38.05%) and courtship transition 9 (89.38 +/− 50.03%).

**Figure 3.**
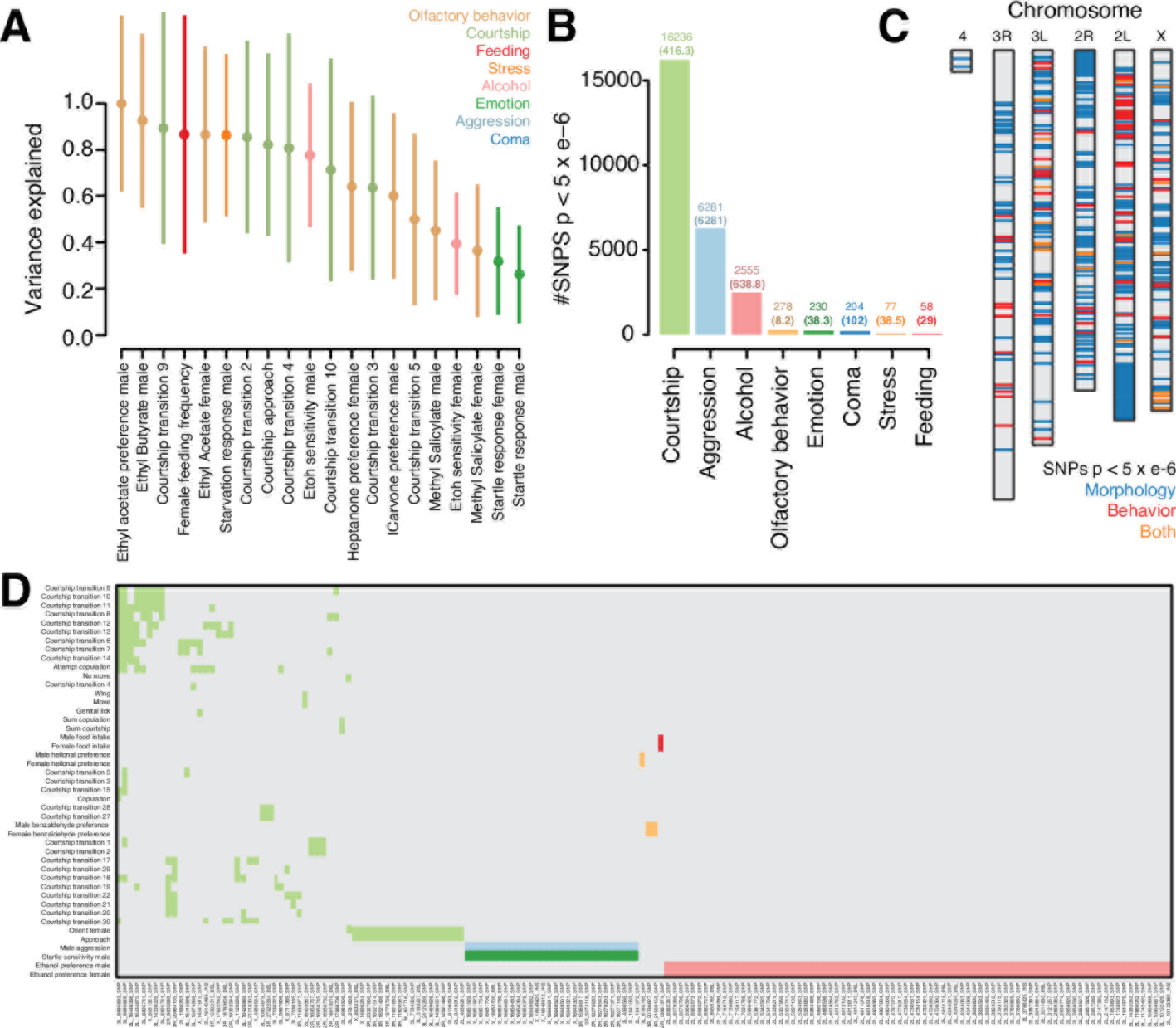
Comparative genome-wide analyses of the Drosophila Genetic Resource Panel. **(A)** Heritability estimates (V(G)/Vp) from GCTA for the 20 measures identified as significant (p-value < 0.05), colored by behavioral category. **(B)** Barplot summarizing the number of SNPs with p < 5 × 10^−6^ collected for each behavioral category from GWAS on 87 traits. **(C)** The distribution of SNPs with p < 5 × 10^−6^ across the *Drosophila melanogaster* genome for morphological (blue) behavioral traits (red) and SNPs that associate with measures of both (orange) **(D)** Heatmap representing the distribution of shared SNPs with p < 5 × 10^−6^ across all behavioral traits. Plotted are SNPs that possess associations with at least two behavioral traits, colored by the categories highlighted in **(A)**.

In addition to an increase in genomic heritability, my QTL analyses also showed that the genomic architectures of courtship and feeding traits may be simpler and of higher effect. To test this I performed a separate GWA experiment for each trait across all lines with available phenotypic data and filtered for SNPs with a nominal p-value of 5 × 10^−6^ (Supplementary methods). At this threshold I found 25,919 SNPs (Fig. 3b; Table S1).

I re-ran GCTA for each trait using only SNPs identified at p < 5 × 10^−6^ from the GWAS (supplementary methods). This test is more conservative compared to genome-wide GCTA since it uses just the fraction of genomic variants significantly associated with each individual trait. After GCTA I found 16 behavioral traits that passed the p-value threshold of P < 0.05. Half of these significant traits were courtship behaviors, including the top four traits with the most variation explained by GWAS SNPs (Fig. S5a). The number of GWAS significant SNPs for these 16 traits varied substantially and was positively correlated with the amount of phenotypic variance explained (Fig. S5b). For traits with more SNPs, significant portions of the variance could be accounted for. For example, 665 SNPs could account for 63.52 +/−8.42% of variation in courtship wing movement, 828 accounted for 68.64 +/−6.69% of genital licking behavior, and 8,013 accounted for 78.45 +/−5.97% of courtship approach behavior. The results from both GCTA tests in the DGRP lines support the hypothesis that courtship and feeding-related behaviors are associated with more heritable genetic architectures of large effect, even within less diverged natural populations.

I next used the DGRP lines to query the extent to which genes or genomic loci may affect multiple behavioral traits (pleiotropy) (Greenspan 2004). I exploited the breadth of phenotypic and genomic data available in the DGRP to empirically address this question at three levels: SNPs, genes, and traits. To allow for comparisons of behavior and other trait types I also conducted GWA for 26 morphological traits reported in Vonesch et al. 2016 (Supplementary methods; Table S2). SNPs found to be associated with morphology and behavior at p < 5 × 10^−6^ were distributed across the *Drosophila melanogaster* genome, 80 of which were associated with both behavioral and morphological traits (Fig. 3c).

With this list of variants I queried which individual SNPs were associated with multiple behavioral categories. I identified 169 SNPs associated with at least two behavioral measures. These variants largely segregated *within* behavioral categories rather than *between* categories, suggesting that at the level of individual SNPs these traits may have largely independent genetic architectures amongst the DGRP lines (Fig. 3d). Many of these SNPs fell within the same genomic regions. I found 72 genes had at least 2 SNPs associated with multiple traits, several of which contained a multitude of variants (Fig. S6a). These genes are enriched for involved in biological processes such as Notch signaling, receptor activity, and morphogenesis (Supplementary methods; Table S3). In addition, I found 81 intergenic SNPs that each occurred within 20kb of their nearest gene - 26 genes in total - suggesting potential regulatory roles for these SNPs (Fig. S6b).

I then assessed the extent to which behaviorally associated variants may act pleiotropically at the trait level, using the list of 25,919 variants associated with behavior. With this I correlated the effect sizes of trait-associated SNPs with the effect sizes of those same variants across all other traits (following ref. 26). The results of this analysis are summarized in the clustered heatmap in Fig. S7. In general I found extensive correlations between behavioral traits, suggesting widespread pleiotropic genetic effects. I also observed several large clusters of highly correlated traits, suggesting a higher-level structure for phenotypic variation based on trait interactions (labeled 1-4 in Fig. S7). The existence of these apparent clusters suggest that, while behavioral categories in the DGRP overlap little in genomic architecture at the individual variant level, there may be common molecular pathways through which different behavioral traits are altered in a correlated fashion.

Finally, I explored pairs of traits with putative directional relationships given the effect sizes of their associated variants. I avoid calling these relationships causal since, given the existence of extensive epistasis and genetic linkage the DGRP lines, it is difficult to identify individual variants of likely causal effect (Huang et al. 2012). I instead sought to elucidate aspects of a directional relationship by discriminating between cases in which a genotype effects multiple traits through different mechanisms versus scenarios where a genotype exerts an effect on a trait through a second, intermediate trait (summarized as *P_1_*←*G*→*P_2_* compared to *G*→*P*_1_→*P*_2_) (Pickrell et al. 2016). In addition to the 87 identified behavioral traits, I included the 26 morphological measures to gather insights into potentially directional relationships between behavior and morphology in the DGRP.

I conducted pairwise tests of each trait at which GWAS variants at the p < 5×10^−6^ level were identified. Using a permutation based test I found 143 trait pairs that showed directionality wherein the correlation of effect sizes was strong and significant in one comparison but not the other (Supplementary methods; Fig. 4a).

**Figure 4.**
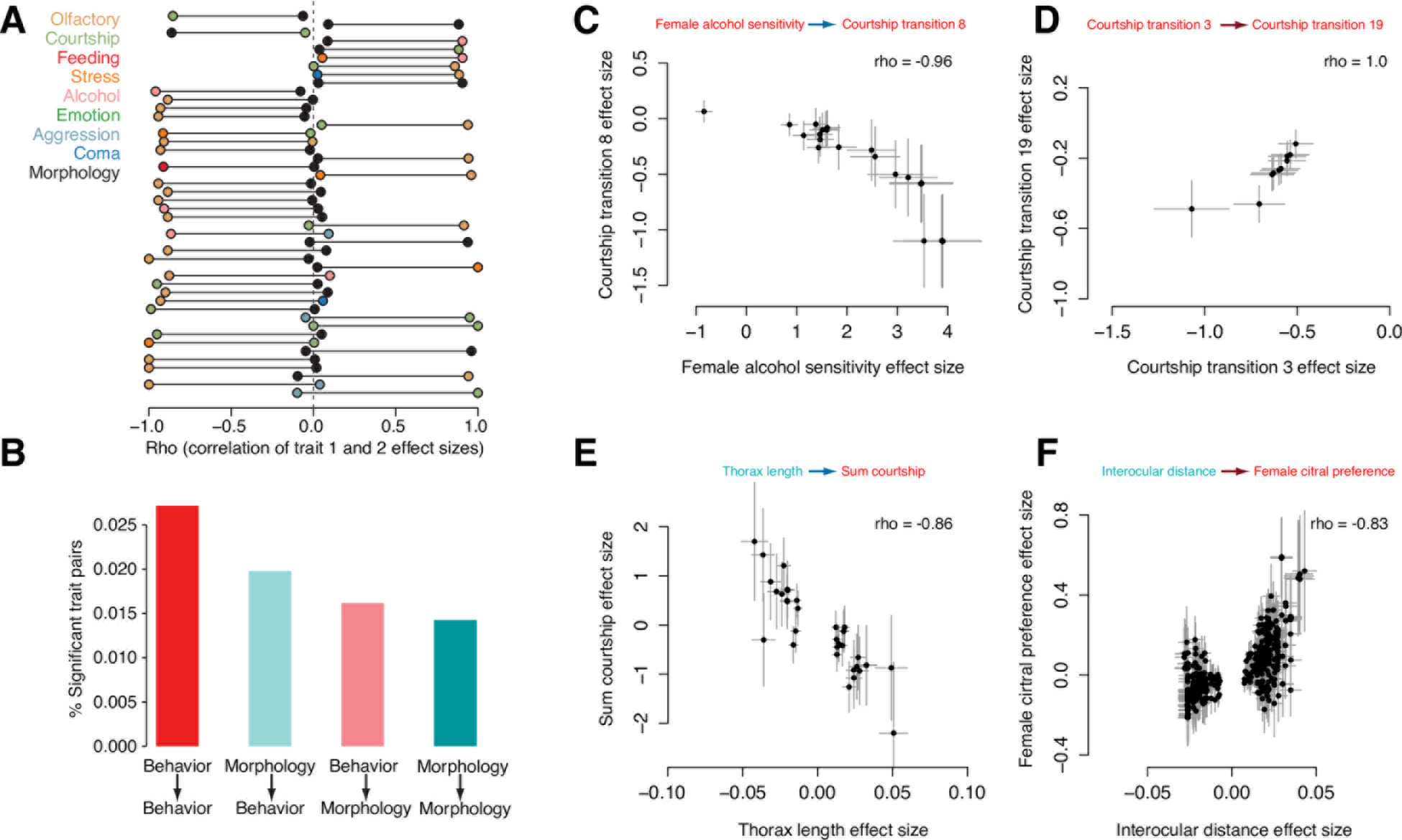
Directional relationships between trait pairs in the DGRP. **(A)** Directional trait pairs identified as significant by permutation testing. Plotted are traits where the significant correlation possesses a rho > 0.85. The significant correlation is represented by a red circle. **(B)** Barplot summarizing the number of significant trait pairs identified where the focal trait is either behavioral or morphological with a correlation one of these two domains. Behavioral focal traits are colored red, morphological traits are colored blue. **(C-F)** Scatterplots of the effect sizes for the focal SNPs of example significant trait ndard errors are plotted as grey lines. Positive correlations are represented by red arrows and correlations are represented by blue arrows.

Trait pairs identified as significant showed an uneven distribution of potential directional effect between behavior and morphology, with the largest amount occurring between pairs of behavioral traits (Fig. 4b). Figures 4c-f highlight examples of these SNP effect size correlations for different behavioral and morphological measures. A particularly interesting connection was found between SNPs associated with EGFR signaling affecting thorax length and the total amount of courtship attempted by male flies (rho=-0.86, p= 8.6×10^−8^; supplementary methods).

The connection between male courtship behaviors and body size has long been recognized in laboratory strains of *Drosophila* though with little evidence of a molecular basis for this effect (Ewing 1961). In general I find extensive evidence of both directional (*G*→*P_1_*→*P_2_*) and general (*P_1_*←*G*→*P_2_*) pleiotropic effects between traits in the DGRP, supporting the notion that the early stages of behavioral diversification involve the role of genes that can effect multiple types of traits. Furthermore, I observe that while variation in behavior across trait categories is associated with non-overlapping variants these may occur in common genes and molecular pathways with pleiotropic effects, reflecting suggestions of the existence of phenotypic “hotspots” that are recurrently used by evolution to sculpt phenotypes (Stern & Orgogozo 2008).

Taken together these results suggest that behavioral traits may respond to evolutionary processes with greater variation than previously appreciated. For example, researchers may now anticipate that assaying a courtship ritual will likely yield a higher genetic effect than, say, variation in a personality trait. These insights are supported by observations that behavioral categories vary in their heritability and genomic architecture during even the earliest stages of diversification within populations. Furthermore, such behaviors are associated with a small number of highly pleiotropic genes and these traits interact, indicating that there are identifiable molecular and phenotypic patterns that govern behavior.

These findings suggest several important caveats and prospects for future behavior genetic studies. First, QTL mapping methods possess inherent limitations in detecting the complete genetic architecture of certain traits. For example, QTL studies are often insensitive to the detection of loci with opposing effects on the trait of interest, thus potentially masking important genetic effects from the researcher’s analysis (Mackay et al. 2009). Future studies of the genetic architecture of behavior will thus benefit from integrating QTL methods with results from genome-wide sequencing and genetic interrogations directed by genome editing^7^. Second, a more complete survey of behavioral categories within and across a variety of taxa are needed to confidently establish whether or not the patterns observed in this study are general principles of how behavior evolves. Finally, empirical tests in the field and lab may offer a deeper understanding of the extent to which courtship and feeding behaviors respond uniquely to selective pressures, and which evolutionary and ecological mechanisms may account for this phenomenon. Expanding on this with the tools and data now becoming available, behavioral biology may begin to produce a more nuanced and predictive understanding of the interplay of genetic forces governing the evolution of behavior.

## Materials and methods

### QTL collection

I first identified behavioral QTL through literature search querying online engines (e.g. PUBMED) with the keywords “QTL”, “behavior”, “quantitative trait locus”, and “behavioral”. I analyzed the results and collected QTL for each relevant publication identified. In order to gather as many relevant QTL as possible over time I expanded the search to include more specific terms relating to behaviors and categories of interest and to those referenced in previously identified papers. I filtered for loci reported as significant by the original authors, resulting in 1,007 QTL from 115 studies. For each locus I recorded the reported effect size (percent phenotypic variation explained), significance measure, genomic location, sample size, and the number of loci reported overall. QTL studies often report other measures in addition to those that I collected (e.g. broad or narrow sense heritability). While it would be desirable to compare certain of these across behaviors and taxonomic groups I found that, within the studies assayed, the reporting of measures other than those I collected was very inconsistent and allowed for only extremely restricted comparisons. Since the measure used to report significance varied across studies I converted all LOD scores to Log p-values using the following R function (R.C.team 2013):

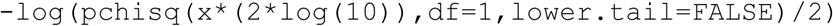

I next classified behaviors following the six groups used in the meta-analysis of mouse QTL studies done by Flint 2003. Several categories represented in our data set were not assayed in this original study (e.g. courtship). In our classification of these I attempted to strike a balance between breadth (to increase the tractability of our comparisons) and biological specificity. To do so I required that a category be represented in at least two species or populations and that the classification match either that reported by the original authors or a reasonable division as reported by the animal behavior literature. The classification of a range of biological traits into broader categories is of course difficult and can repeatedly tempt debate; accordingly this is discussed at length in Flint 2003. I offer that it is important to rigorously test results implicating a broadly defined category as interesting through comparisons of that category to the overall distribution of effects, with the goal of controlling for bias introduced by the original classifications (as is discussed below). All QTL and the associated measures mentioned here are available in Table S1.

### Phylogeny

I used the phylogenetic relationships reported in Ponting 2008 as a template for our phylogeny of species examined (Fig. 1). I added unrepresented species and adjusted dates of evolutionary divergence using the most recent reports available for each specific clade/species. The following sources were used (along with the associated phylogenetic divergences):

*Ruff/quail and chicken*: Jarvis et al. 2014
*Quail and chicken*: Kayang et al. 2006
*Nine spined and three spined stickleback*: Guo et al. 2013
*Stickleback and teleost*: Pfister et al. 2007
*Cave fish and teleost divergence*: Briggs 2005
*Laupala cricket and insect divergence*: Misof et al. 2014
*Wax moth and insects*: Misof et al. 2014
*Pea aphids and insects*: Misof et al. 2014
*Peromyscus and mice/rats*: Bedford and Hoekstra 2015
Solenopsis *and* Apis: Ward 2014
*Sheep and cows*: Bibi 2013
*White fish and teleosts*: Betancu-R et al. 2013

### Effect size comparisons

The overall distribution of effect sizes (Fig. 1B) was plotted using the density function in R. Since some behavioral categories possessed relatively small sample sizes all comparisons of effect size were done with the non-parametric Kruskal-Wallis test.

For the analyses plotted in Fig. 2a-2c and Figs. S1-4 I summed the effect sizes of all loci associated with a specific behavioral measure for each study. This was done to allow for a comparison of the maximum amount of phenotypic variance explained for each trait in order to allow for conservative test between courtship and feeding and all other traits. For example, there may have been non-courtship/feeding traits associated with many loci that, on their own, possessed small effects but when added together explained a substantial portion of variation. Following this I filtered for loci where sample size information and evolutionary divergence information were available, resulting in 773 loci. The rationales for each estimate of evolutionary divergence are discussed in the next section.

I used several linear models to test for potential biasing effects from evolutionary divergence and sample size, both individually and combined. The resulting residuals from these models are presented in figures S1-3. I found no significant correlations between the residuals from the models and the original variables tested. For the model incorporating sample size alone I observed a correlation between the residuals and sample size itself of −4.06 × 10^−17^. For the models incorporating evolutionary divergence alone I observed a correlation between the residuals and divergence of −5.34 × 10^−17^. Finally for the model incorporating both there was a correlation of −1.16 × 10^−16^ with sample size and a correlation of 3.00 × 10^−17^ for divergence. Given the lack of correlation with either variable this suggests that the combined model successfully controlled for both factors. The residuals for this final combined model were then used for a comparison between all categories (Fig. S1) and between courtship/non-courtship (Fig. 2a) and feeding/non-feeding (Fig. S4a).

For the comparison of the observed courtship and feeding residual effect sizes to the quantiles of all non-courtship/feeding traits I used the following R function (where `non_courtship` and `courtship` are vectors of residual effect sizes for these groups):

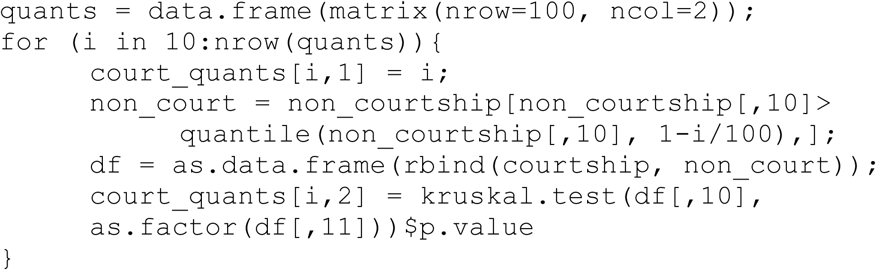

The bootstrap comparisons in Figs. 2c and S3c were done using the custom R function `bootstrap.2independent` which is available on the Fernald lab website. For these tests I permuted the non-courtship/non-feeding residual effect sizes 10,000 times (with replacement) to create a null distribution against I which I tested the observed median residual effect size for each trait. A p-value for each test was calculated by dividing the sum of instances in which the permuted medians were greater than the observed by 10,000. All plots were produced using base graphics in R and adjusted for design in Adobe Illustrator.

### Data collection of the DGRP lines

I downloaded the DGRP freeze 2.0 variant calls and plink files from the *Drosophila* genetics reference panel website (http://dgrp2.gnets.ncsu.edu). Raw data for phenotypic measures were downloaded from the following sources:

*Starvation resistance*: Mackay et al. 2012
*Startle response:* Mackay et al. 2012
*Chill coma recovery time*: Mackay et al. 2012
*Startle response under oxidative stress*: Jordan et al. 2012
*Negative geotaxis under oxidative stress*: Jordan et al. 2012
*Olfactory behavior (benzaldehyde):* Swarup et al. 2013
*Courtship behavior*: Gaertner et al. 2015
*Olfactory behavior (multiple measures):* Arya et al. 2015
*Aggressive behavior*: Shorter et al. 2015
*Food intake*: Garlapow et al. 2015
*Alcohol sensitivity*: Morozova et al. 2015
*Morphology*: Vonesch et al. 2016

I compiled the raw data into two tables for use in genome-wide analyses of SNP variation, one composed of the 87 behavioral traits obtained and another of the 26 morphological traits. For traits in which multiple measurements were reported I calculated the mean trait measurement and used this for subsequent analyses. I classified traits into behavioral categories in the same fashion as for the evolutionary QTL analyses.

### Heritability analyses

I first employed genome-wide complex trait analysis (GCTA) to survey genomic heritability across the 87 behavioral traits (Yang et al. 2011). For each trait I used `greml v1.26.0` to obtain estimates of heritability from genome-wide SNP variation across all DGRP lines for which phenotypic measures were available. Using the plink files obtained from the DGRP website (base file name `dgrp2`) I first created a genotype relatedness matrix for all DGRP lines:

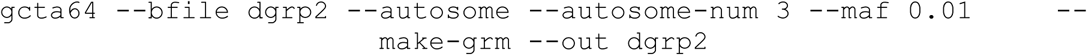

Inidividual phenotype files (`*.phen`) were created for each trait, including fam and individual IDs and the associate phenotypic measures for each DGRP line. I ran `GREML` for each phenotype separately:

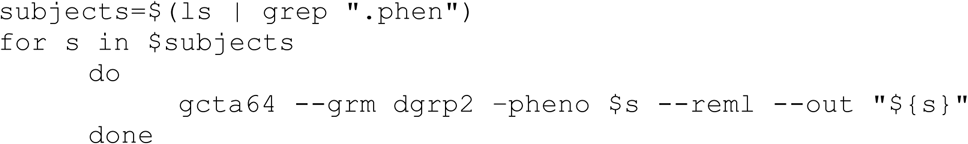

I then filtered for traits in which the reported p-value from `GREML` was <0.05, resulting in 20 traits. Fig 3a. shows the distribution of phenotypic variance explained by genome-wide SNPs as measured by the genotypic variance divided by phenotypic variance (Vg/Vp).

For the GCTA analyses of just GWAS significant SNPs I compiled a list of associated SNPs for each trait and built a separate genotype relatedness matrix for each by extracting just those SNPs from the plink bed files. I then reran `GREML` for each trait using the corresponding genotype relatedness matrix and testing only for the SNPs that it contained. Like above I then filtered for traits in which the reported p-value from `GREML` was <0.05, resulting in 16 traits.

### Genome-wide association analyses

The plink and phenotype files from the GCTA analyses were used to conduct separate genome-wide association studies (GWAS) for each trait. I used `plink v1.90` to conduct these tests on the combined phenotype matrix (`“dgrp_phenos.txt”`):

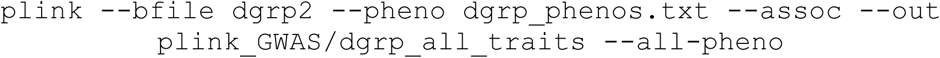

Associations were then filtered for a p-value < 5 × 10^−6^. SNPs associated with multiple traits were identified and plotted using a binary heatmap with the `heatmap2` function in R. Genes associated with multiple SNPs were identified using the variant annotation file available on the DGRP website.

I next assayed relationships between SNPs and multiple traits using the effect sizes (betas) in the `*.qassoc` files outputted by `plink`. To do so I compiled a matrix of the effect sizes for all traits at each of the 25,919 significant SNPs (Table S4). This matrix could then be directly queried for comparison of the effect sizes associated with a certain set of SNPs across traits of interest. In order to assess the overall structure of this data set I used Spearman rank correlations to test the associations between all possible trait pairs. The results of this test were visualized using the clustering functionality of `heatmap2` in R (Fig. S7).

### Tests for trait pair directionality

Directionality in the relationships between trait pairs was tested by first obtaining pairwise rank correlations for each trait pair in which both traits were associated with >3 significant SNPs (60 traits). For traits x and y, s_1_ is the vector of SNPs significantly associated with trait x and s_2_ is the vector of SNPs significantly associated with trait y. `xx` is the vector of effect sizes at s^1^ for trait x and `xy` is the vector of effect sizes at s_1_ for trait y. Similarily `yy` is the vector of effect sizes at s_2_ for trait y and `yx` is the vector of effect sizes at s_2_ for trait x. Rank correlations can then be obtained for each in R:

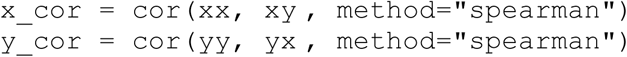

Since the strongest signal of directionality would be cases in which the absolute value of `x_cor/y_cor` equals 1 and the other is equal 0, I assessed directionality as a function of how close to 1 the absolute difference between the correlations was:

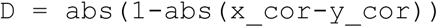

I filtered for trait pairs in which rho for one correlation was >0.5 and for the other was <0.1. I then tested the directional significance of each trait pair by permuting `xx`, `xy`, `yy`, and `yx` 1,000 times and recomputed `x_cor, y_cor`, and `D` for each permutation. I then calculated a p-value for each trait pair by comparing the vector of permuted `D` values (`pseudo`) to the observed `D`:

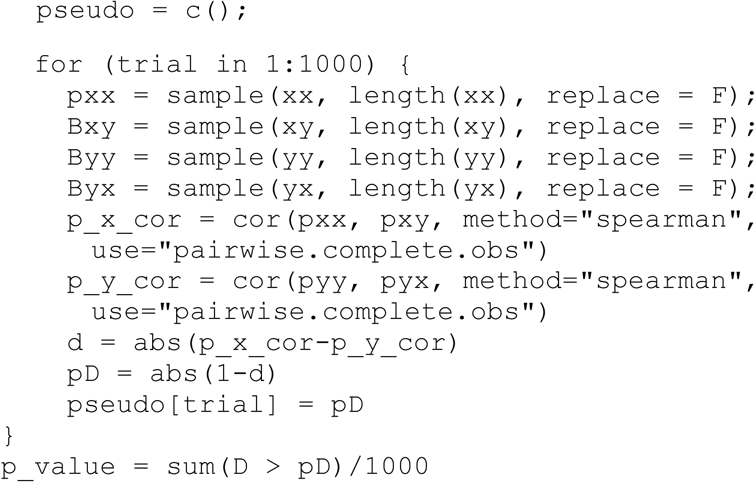

The resulting p-values were adjusted using Bonferroni correction.

## Acknowledgments

I thank Hunter Fraser, Russell Fernald, Christopher Martin, Graham Coop, David Kingsley, Austin Hilliard, Trudy Mackay, and David Stern for critical reading and useful discussions.

